# The ancient cardioprotective mechanisms of ACE2 bestow SARS-CoV-2 with a wide host range

**DOI:** 10.1101/2021.01.03.425115

**Authors:** Gianni M. Castiglione, Lingli Zhou, Zhenhua Xu, Zachary Neiman, Chien-Fu Hung, Elia J. Duh

## Abstract

SARS-CoV-2 infects a broader range of mammalian species than previously anticipated, suggesting there may be additional unknown hosts wherein the virus can evolve and potentially circumvent effective vaccines. We find that SARS-CoV-2 gains a wide host range by binding ACE2 sites essential for ACE2 carboxypeptidase activity. Six mutations found only in rodent species immune to SARS-CoV-2 are sufficient to abolish viral binding to human and dog ACE2. This is achieved through context-dependent mutational effects (intramolecular epistasis) conserved despite ACE2 sequence divergence between species. Across mammals, this epistasis generates sequence-function diversity, but through structures all bound by SARS-CoV-2. Mutational trajectories to the mouse conformation not bound by SARS-CoV-2 are blocked, by single mutations functionally deleterious in isolation, but compensatory in combination, explaining why human polymorphisms at these sites are virtually non-existent. Closed to humans, this path was opened to rodents *via* permissive cardiovascular phenotypes and ancient increases to ACE2 activity, serendipitously granting SARS-CoV-2 immunity. This reveals how ancient evolutionary trajectories are linked with unprecedented phenotypes such as COVID-19 and suggests extreme caution should be taken to monitor and prevent emerging animal reservoirs of SARS-CoV-2.

**One sentence summary:** A conserved mechanism essential for ACE2 catalytic activity is exploited by SARS-CoV-2 binding, allowing the virus to infect a wide range of species.

## Introduction

SARS-coronavirus-2 (SARS-CoV-2) is thought to have evolved from an ancestor (RaTG13) isolated from the Chinese horseshoe bat (*Rhinolophus sinicus*) (*1*), potentially fusing with viruses from an intermediate source (e.g. pangolin; *Manis javanica*) (*2*). These zoonotic origins ultimately led to the evolution of a virus that is highly transmissible among humans, causing an unprecedented public health emergency (*3*, *4*). A wide phylogenetic range of mammalian species have been demonstrated to be susceptible to SARS-CoV-2 infection, including non-human primates, dogs, cats, ferrets, hamsters and minks (*5*–*12*). Characterizing the entire host range of SARS-CoV-2 is important for identifying the risks of anthroponosis and zoonosis between humans and other species, which can pose a major health risk by forming novel viral reservoirs where new mutations can evolve (*13*, *14*). A recent example is mink farms, where infection by humans led to the evolution of novel viral strains, which have since reinfected human populations (*11*). Attempts to predict the host range of SARS-CoV-2 have greatly underestimated the extent to which SARS-CoV-2 can infect certain non-primate species, especially non-felid carnivores including mink, dog and ferret (*5*–*11*, *15*). These predictive methods depend on comparative sequence analyses of angiotensin-converting enzyme 2 (ACE2) – the cellular receptor for SARS-CoV-2 – and scoring based on sequence and structural homology to the human ACE2 viral binding interface (*15*–*11*). The difficulty of predicting the SARS-CoV-2 host range through these methods proves that the virus can bind a wide-range of ACE2 orthologs despite extensive sequence divergence. This opens the possibility for the existence of additional unknown mammalian hosts.

Infection by SARS-CoV-2 is mediated by the binding of the viral S-protein receptor binding domain (RBD) to ACE2 (*1*, *18*), displaying a nanomolar affinity higher than that of SARS-CoV-1 (*19*). Previously, it was identified that a single mutation to ACE2 was sufficient to decrease SARS-CoV-1 RBD binding by nearly 90% (*20*). By contrast, structural analyses have identified that the SARS-CoV-2 RBD targets multiple binding ‘hotspots’ within the ACE2 ectodomain, presenting a diffuse and multifaceted binding strategy (*21*–*23*). While many sites within the viral binding domain of ACE2 can modulate binding of the SARS-CoV-2 S protein RBD (*24*), it remains unclear how amino acid interactions between these sites facilitate viral binding and infection. The ability of the SARS-CoV-2 S protein RBD to bind divergent sequences may be therefore be explained by ACE2 intramolecular epistasis, a form of context dependence, where amino acids have different functional effects depending on residues at other sites (*25*). Although intramolecular epistasis can limit the number of possible amino acids combinations that will confer a given function (thus making evolution ‘predictable’; (*26*, *27*)), it can also give rise to novel compensatory interactions following an ‘original’ mutation (*28*–*31*), ultimately generating completely different amino acid combinations converging on the same structural/functional ‘solution’, making sequence-function evolution unpredictable (*32*–*36*). If pervasive intramolecular epistasis exists within the viral binding interface of ACE2, then this may explain why ACE2 sequence homology alone has not been a reliable predictor of viral binding and infection.

In mammals, ACE2 evolved to serve an essential reno- and cardio-protective role *in vivo*, mediated through its regulation of the renin-angiotensin-aldosterone system (RAAS) in conjunction with ACE. The signaling peptide Angiotensin-II (Ang-II) generated by ACE—a major clinical target for hypertension—stimulates vasoconstriction, inflammation, and fibrosis responses through the Ang-II/AT_1_R axis (*37*, *38*). ACE2 carboxypeptidase activity counteracts these effects through conversion of Ang-II to Ang-(1-7), a peptide which induces vasodilatory, anti-inflammatory, and anti-fibrotic effects through MAS signaling (*38*, *39*). Loss of ACE2 in mice worsens cardiac dysfunction in obesity, increases diabetic kidney dysfunction, increases mortality rates after myocardial infarction, and can have severe effects on cardiac contractility (*37*, *39*–*41*). This protective role of ACE2 is largely mediated through its enzymatic processing of Ang II to Ang-(1-7), where exogenous delivery of either ACE2 or Ang-(1-7) can protect against pathogenic features of multiple cardiovascular and kidney diseases (*37*, *41*, *42*). Given the myriad protective roles of ACE2 enzymatic activity, it may be expected that ACE2 function is highly conserved across species. However, mice display ~ 50% higher ACE2 activity relative to humans (*43*), and are insensitive to the vasodilatory effects of Ang-(1-7) (*39*), suggesting increased ACE2 activity in rodents may have evolved to serve non-vasodilatory protective functions. These observations suggest that minor differences in ACE2 function can drive major physiological differences between species, as seen in the evolution of other protein systems (*27*, *31*, *44*). Organismal sensitivity to mutations affecting ACE2 activity may be especially pronounced due to the X-chromosome location of the *ACE2* gene (*45*), where even heterozygous *ACE2* knockout females (-/x) display increased susceptibility to heart and kidney injury (*39*).

If natural variation across species has evolved to shape ACE2 function, then the sequence recognized by SARS-CoV-2 may have a pleiotropic role in mediating ACE2 function. We therefore investigated whether SARS-CoV-2 binding to ACE2 could be abolished *without* disrupting ACE2 enzyme function. To expedite this, we took an evolutionary approach that leveraged the natural sequence variation found in mouse ACE2, which SARS-CoV-2 is unable to bind (*1*, *18*). Here we identify a specific combination of mutations unique to rodents which fully abolishes RBD binding when inserted into human and dog ACE2, but which in isolation, significantly decreases ACE2 enzyme activity. These detrimental intermediates would likely severely compromise the cardio- and reno-protective functions of ACE2 activity, explaining why these mutations are rare across mammalian species.

## Results

We investigated ACE2-RBD binding as well as ACE2 enzymatic function across a range of boreoeutherian mammals either susceptible or immune to SARS-CoV-2 infection [human (*Homo sapiens*), dog (*Canis lupus familiaris*), and pangolin (*Manis javanica*); *vs*. mouse, (*Mus musculus*) and Chinese horseshoe bat (*Rhinolophus sinicus*), respectively] (*1*, *7*, *9*, *18*). Using flow cytometry, we found a trend of significantly stronger association of the SARS-CoV-2 RBD with both human and pangolin ACE2 relative to that of CoV-1, consistent with previous studies (*19*) (Fig. 1A-D). Notably, CoV-1 and −2 RBD association was strongest with human ACE2. We also found evidence that the RBD of both SARS-CoV-1 and −2 S protein could not bind mouse nor bat ACE2 (Fig. 1B-D), consistent with previous studies and reaffirming pangolin as a likely intermediate between bat and human (*2*, *7*, *9*, *18*, *46*). To complement these binding assays, we next characterized the ability of ACE2 orthologs to enable SARS-CoV-2 S pseudovirus entry into cells. We transfected ACE2 orthologs into human cells (HEK293T) and exposed these cells to pseudotyped MLV particles containing the SARS-CoV-2 S protein. Consistent with our flow cytometry data and with previous studies, this assay recapitulated the host range of WT SARS-CoV-2 (*18*), displaying significant infection of cells expressing human, dog, and pangolin ACE2, but not that of mouse or bat (Fig. 1E). To investigate whether this variation in ACE2-SARS-CoV-2 S binding is mirrored by functional variation in ACE2 enzyme activity, we measured the carboxypeptidase activity of ACE2 orthologs *in vitro* using a well-characterized fluorometric biochemical assay (*43*) (Materials and Methods; Fig. S1). Strikingly, we found significant variation in ACE2 hydrolysis rates across all mammalian species, suggesting tuning of enzymatic function across evolutionary history (Fig. 1F-G), perhaps in response to interspecies RAAS variation. Interesting, dog, bat and pangolin all displayed low ACE2 hydrolysis relative to human and mouse. Mice displayed the highest ACE2 hydrolysis rates, consistent with previous reports (*43*). This demonstrates that ACE2 function varies considerably across species, and suggests that ACE2 sequence variation may reflect diversification of ACE2 enzymatic function.

**Fig 1.**
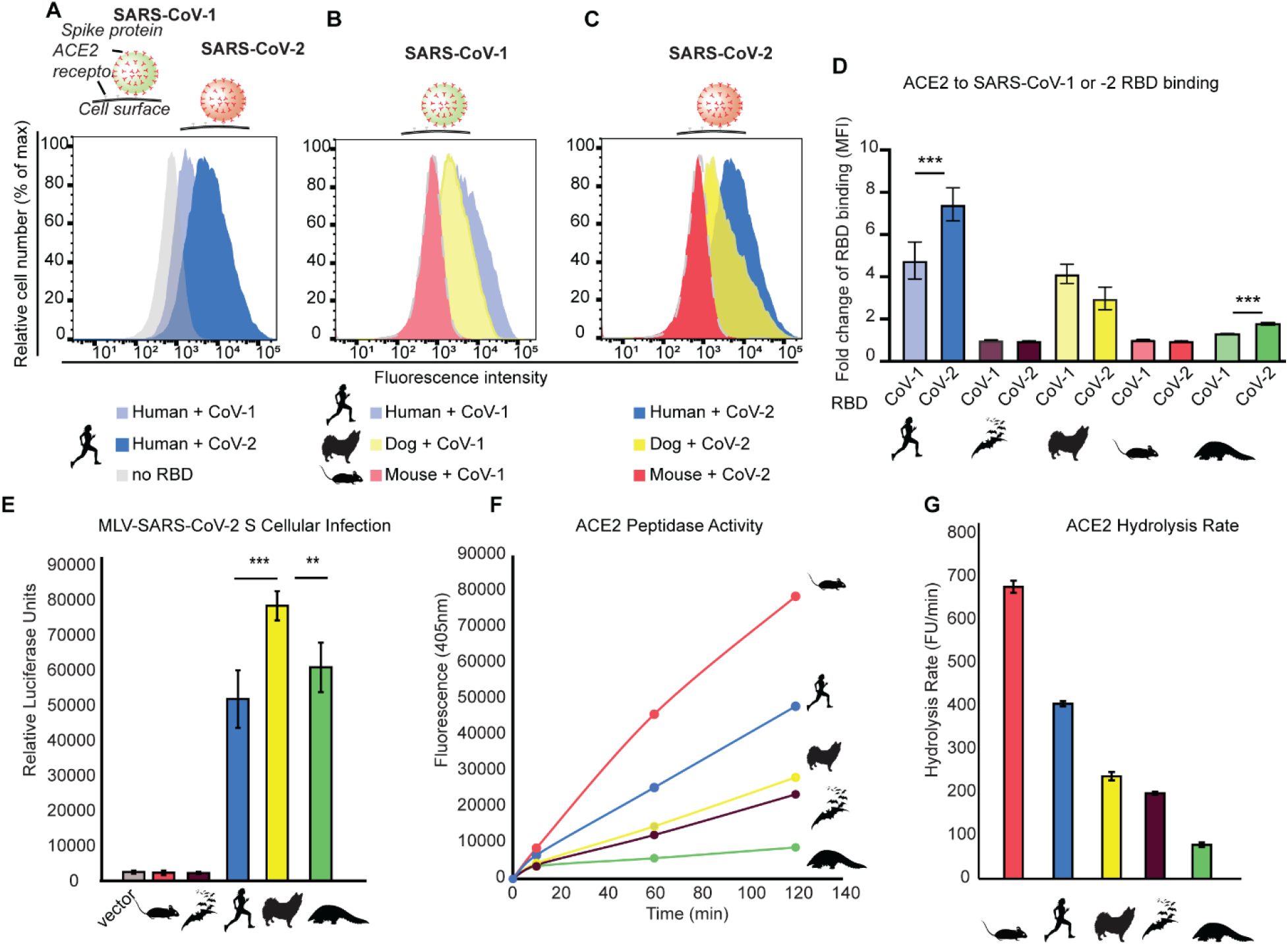
Natural variation in SARS-CoV-2 binding is mirrored by diversity in ACE2 enzyme activities. (A-D) Flow cytometry was used to quantify SARS-CoV-1 and CoV-2 RBD-Fc association with human cells (HEK293T) expressing ACE2-eGFP orthologs from various mammalian species (mean fluorescence intensity; MFI (*19*)). (E) SARS-CoV-2 S pseudovirus infection of HEK293T cells expressing ACE2 orthologs was quantified using a luciferase reporter system. (F) The carboxypeptidase activity of ACE2 was quantified using a fluorometric peptide incubated with solubilized HEK293T cells transfected with ACE2 orthologs. (G) Hydrolysis rate of ACE2 orthologs (Fluorescence units/minute).

To test this, we first took a bioinformatic approach. We reasoned that since ACE2 has evolved under pressures related to its enzymatic processing of Ang-II/Ang-(1-7), signatures of natural selection at sites within the viral binding interface may reflect a potential role of those sites in mediating ACE2 catalytic activity (*47*). We therefore searched for shifts in *ACE2* mutational rates (d_*N*_/d_*S*_) beyond what may be expected from neutral evolutionary pressures alone (*48*, *49*). To do this, we constructed a phylogenetic dataset representing full-length mammalian *ACE2* sequences (Fig. S2; Table S1) and conducted a codon-based statistical phylogenetic analysis (PAML, HyPhy). We found significant evidence that mammalian ACE2 is under positive selection, outperforming models assuming neutral molecular evolution (Table S2). This provides evidence that mammalian ACE2 functional diversification is a product of natural selection, rather than neutral processes.

To focus our analysis on ACE2 sites responsible for mediating SARS-CoV-2 binding, we took an evolutionary approach leveraging natural sequence variation found in mouse ACE2, which SARS-CoV-2 is unable to bind (*1*, *18*). We identified a set of six residues unique to mouse ACE2 that were of particular interest due to their proximity to viral RBD residues implicated in the ACE2-RBD structure (Fig. 2A). These sites contact opposite ends of the viral RBD, facilitating binding through a combination of hydrogen bonds (Q24, K353), as well as van der Waals forces within a hydrophobic pocket of ACE2 (L79, M82, Y83, P84) (*21*, *23*). These ACE2 residues are located outside the ACE2 active site mediating catalysis of Ang-II/Ang-(1-7) (Fig. 2A; yellow), suggesting that any role of these sites in mediating ACE2 function would likely be through indirect effects modulating the protein structure, as seen in other protein systems (*47*, *50*). Through phylogenetic methods (Table S2; M8; Bayes empirical Bayes), we found that several of these sites deviated significantly from neutral expectations (sites 24, 79, dN/dS>1; site 353, dN/dS<1), suggesting potential functional roles (Fig.2B). Interestingly, these sites are less variable in primates relative to other mammalian lineages (Fig. 2B; (*16*)), displaying significantly decreased *ACE2* evolutionary rates relative to bats (Chiroptera) and rodents (Table S3; (CmD; dN/dS). These evolutionary signatures suggest that the RBD binding interface of ACE2 may play a role in the functional diversification of ACE2 catalytic activity, with extant residues in primates being strongly constrained.

**Fig 2.**
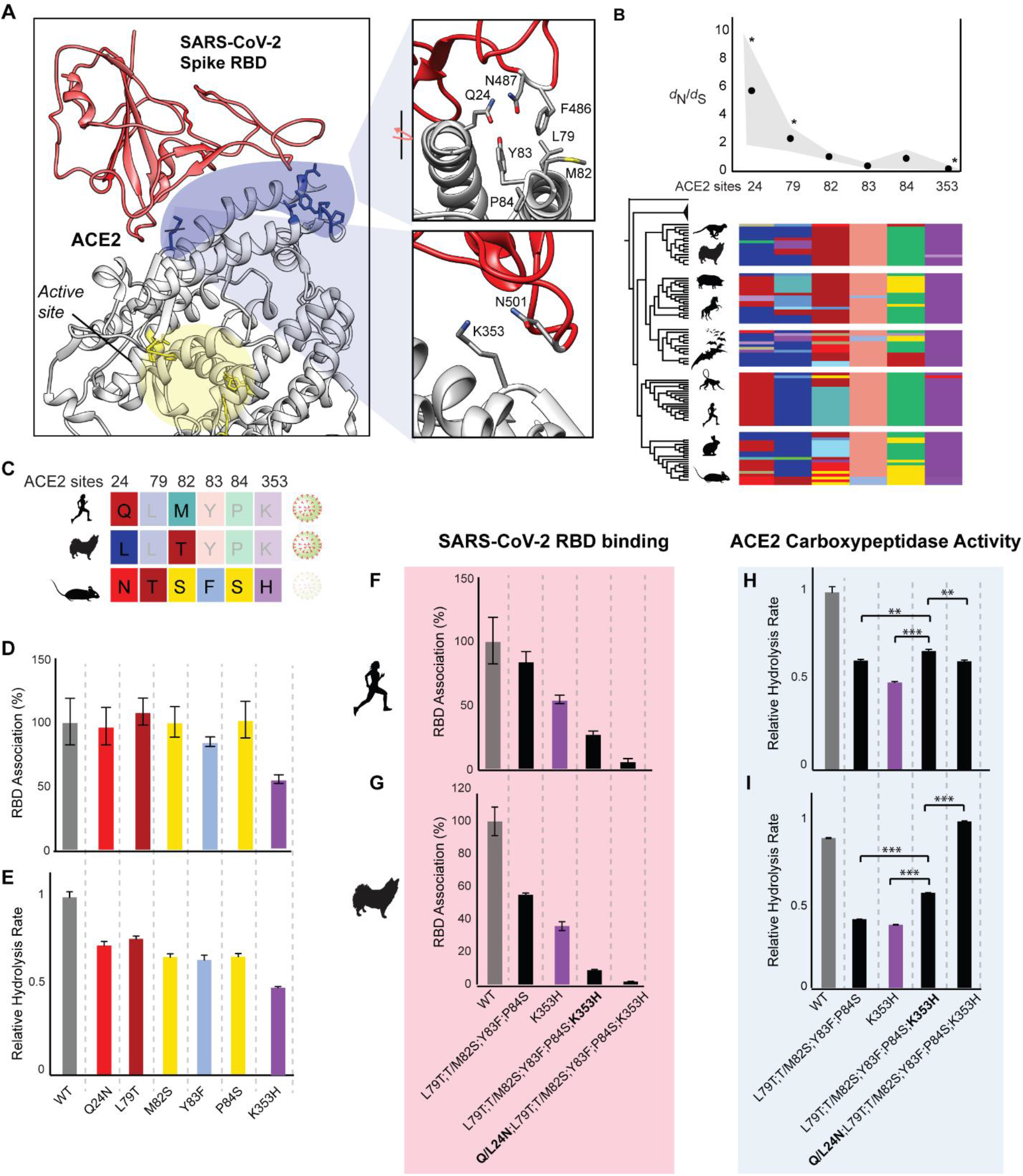
Amino acid interactions mediating ACE2 activity govern SARS-CoV-2 binding to human and dog ACE2. (**A**) SARS-CoV-2 gains cellular entry through the viral spike protein receptor binding domain (RBD, red), which targets binding hotspots on the ACE2 receptor (blue) distal to the ACE2 active site (yellow) [6M17; (*23*)]. (**B**) Statistical phylogenetic analyses (dN/dS averages [dots] and ranges [grey]; PAML, HyPhy) of an alignment of mammalian ACE2 coding sequences reveals positive and negative selection (*) on RBD binding hotspots. Alignment of ACE2 residues across boreoeutherian mammals is shown. (**C)** SARS-CoV-2 is unable to bind mouse ACE2, which displays unique amino acid residues at positions within the RBD binding interface relative to other mammals. (**D**) Flow cytometry was used to quantify RBD association with human cells (HEK293T) expressing WT and mutant ACE2. (**E**) The effect of ACE2 mutations on ACE2 hydrolysis rates was measured using a fluorometric peptide (**F-G**) Interaction effects between amino acids on opposite ends of the viral binding interface determine RBD binding (F-G) and ACE2 activity (H-I) in both human (F, H) and dog ACE2 (G, I).

We next experimentally characterized the role of these sites on both SARS-CoV-1 and CoV-2 binding and infection, as well as their effect on ACE2 catalytic activity. We did this by substituting the rare residues found in mouse ACE2 into human and dog ACE2 (Fig. 2C). First, we characterized the role of single mutations on the binding of the SARS-CoV-1/2 S RBD to human WT and mutant ACE2 using co-immunoprecipitation and flow cytometry. In contrast with SARS-CoV-1 (Fig. S3), a single mutation was not sufficient to abolish SARS-CoV-2 RBD binding to ACE2 (K353H; Fig. 2D). Indeed, most single mutations had only minor effects on SARS-CoV-2 RBD binding (Fig. 2D). However, each of the six single mutations displayed considerable effects on human ACE2 hydrolysis activity (Fig 2E). This demonstrates that ACE2 residues within the RBD binding interface can indirectly modulate ACE2 activity.

Next, we attempted to abolish RBD binding to both human and dog ACE2 by making multiple substitutions at these ACE2 sites. Strikingly, RBD binding to the ACE2 of both species was abolished with just these six substitutions (Fig. 2F-G), despite numerous other sites implicated in mediating interactions with the viral RBD (*21*, *23*). Using the MLV-SARS-CoV-2 pseudovirus system, we found that these six substitutions in combination also significantly reduced pseudovirus infection of HEK293T cells expressing wild type and mutant human and dog ACE2 (Fig. S4). In both species, abolition of RBD binding depended on mutating sites 24 and 82 (Fig. 2F-G), despite displaying different residues in human (Q24; M82) and dog (L24; T82) ACE2 (Fig. 2C). This demonstrates that SARS-CoV-2 utilizes the same combination of ACE2 positions to bind both human and dog ACE2 despite amino acid variation at these sites. This may explain why bioinformatic predictions of SARS-CoV-2 host range based on human ACE2 sequence homology have tended to underestimate the infection risk of species such as dogs, ferrets, and minks (*6*, *7*, *9*–*11*, *15*).

We reasoned that this could be due to conserved compensatory interactions between ACE2 sites, allowing different residues to ultimately maintain the ACE2 structure bound by SARS-CoV-2. Consistent with this, we observed in both human and dog ACE2 a context-dependence (epistasis) of mutational effects between the distal domains of the RBD binding interface. Specifically, the quadruple mutation (L79T; T/M82S; Y83F; P84S) had a much larger effect on RBD binding when a single mutation on the opposite side of the ACE2 binding cleft were included the genetic background (K353H) (Fig. 2F-G). Epistasis between these distal sites indicates that residues at one end of the ACE2-RBD binding cleft can modulate the structural effects of residues on the other end. We hypothesized that the specific amino acid combinations at these sites therefore originally evolved to maintain the ACE2 structure underpinning ACE2 catalytic activity. Consistent with this, we found that the detrimental functional effects of the quadruple mutant (L79T; T/M82S; Y83F; P84S) as well as the K353H mutant were significantly reversed when combined—a phenomenon known as sign epistasis—displaying partial compensation for each other’s detrimental effects on ACE2 hydrolysis rates (Fig. H-I). Further evidence of epistasis in this ACE2 domain is seen in the sextuple mutant, where introduction of L24N in dog ACE2 fully rescued ACE2 activity, whereas Q24N decreased activity in human relative to the quintuple mutant (Fig. H-I). This discrepancy is likely attributable to amino acid interactions between site 24 and other sites not investigated. Taken together, these results demonstrate that extensive amino acid interactions within the RBD binding interface of ACE2 indirectly mediate ACE2 enzymatic activity. This is achieved through compensatory interactions that modulate the ACE2 structure recognized by SARS-CoV-2. This mechanism is conserved in dog and human ACE2, which are separated by ~90 million years of evolution (*51*), suggesting these structural interactions are a general feature of mammalian ACE2 evolution. Thus, compensatory interactions in this domain appear to have facilitated divergent amino acid combinations to converge and generate the same structure recognized by SARS-CoV-2 (*32*–*36*). Hence, our evidence strongly suggests that the unpredictable and wide host range of SARS-CoV-2 is likely due to this ancient epistasis of ACE2.

Why did these substitutions serendipitously blocking SARS-CoV-2 binding only evolve in some rodent species? The compensatory interactions inherent to intramolecular epistasis can both open and close evolutionary trajectories depending on how accessible favorable mutational combinations are within evolutionary space (*31*, *52*–*54*). In humans and dogs, this trajectory appears to have been closed by the presence of deep valleys containing detrimental intermediates with compromised ACE2 function (Fig. 3A). Traversal of evolutionary valleys is thought to be associated with fluctuations in environmental variables, which can change physiological constraints on protein function (*31*, *53*). Since ACE2 activity plays major roles in the cardiovascular system, we reasoned that cardiovascular differences between mammalian species may result in differential constraints on ACE2 sequence-function. A central cardiovascular role mediated by ACE2 activity is the regulation of blood pressure *via* enzymatic processing of ANG- (II) to ANG-(1-7) (Fig. 3B). We found extremely low human allele frequencies for missense polymorphisms at these sites (gnomAD), with most at zero (Fig.3C). This is consistent with the role of these sites in mediating ACE2 activity and with the known association of human ACE2 polymorphisms with hypertension (*55*, *56*). We reasoned that these physiological constraints would be more relaxed in animals with low blood pressures, permitting sequence-function evolution. We conducted a phylogenetic statistical analysis on systolic blood pressure values across mammalian species and observed a significant correlation with body size (r^2^=0.26; p=0.001; phylogenetic independent contrast least squares linear regression) (Fig. 3D; Fig. S6). Blood pressures were lowest in rodent and bat species—two lineages immune to SARS-CoV-2 (*18*, *46*) which also display the highest *ACE2* evolutionary rates relative to other mammals (aBsREL; Fig. 3E; Table S2). This is consistent with bat (*Rhinolophus sinicus*) ACE2 displaying the lowest hydrolysis rates relative to all other species investigated here (except Pangolin, for which blood pressure measurements are not available to our knowledge). Interestingly, rodents are insensitive to the vasodilatory effects of Ang-(1-7) (*39*), suggesting their high ACE2 activity may be unrelated to low blood pressure. Although speculative, relaxed physiological constraints on the vasodilatory role of ACE2 function (due to small body size) may explain why rodents were situated to tolerate an evolutionary trajectory that presumably contained detrimental intermediates as seen in dog and human ACE2 mutants.

**Fig 3.**
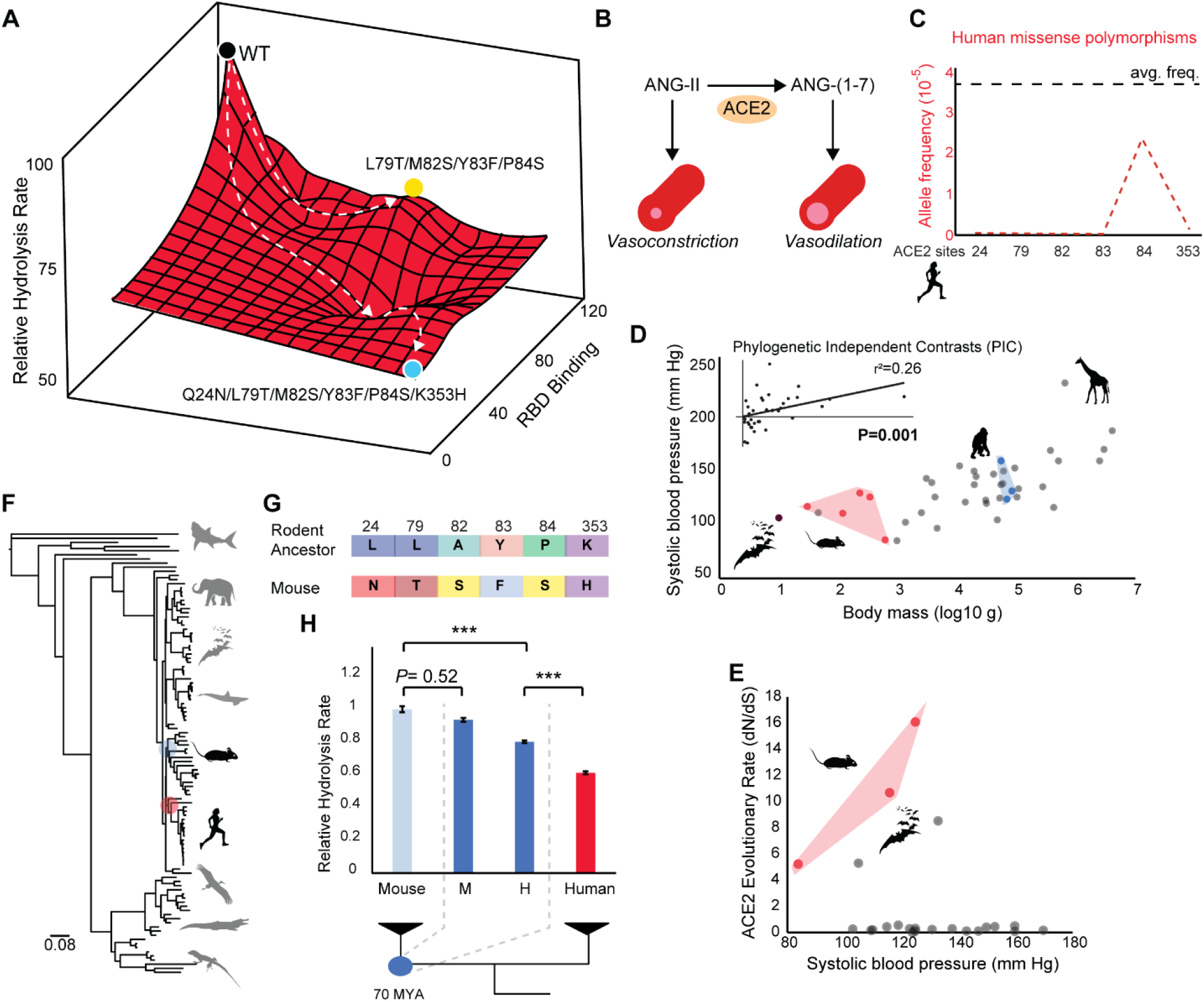
Human polymorphisms required for SARS-CoV-2 immunity are blocked by evolutionary valleys and cardiovascular constraints. (**A**) Substitution of mouse residues into human ACE2 abolishes ACE2-RBD binding, but with large trade-offs on ACE2 hydrolytic activity. Sign-epistasis in human and dog ACE2 activity creates a jagged landscape of intermediate valleys that block this evolutionary trajectory. (**B**) Hydrolysis of ANG-II by ACE2 generates ANG-(1-7), a vasodilatory peptide (*39*). (**C**) Allele frequencies for human missense polymorphisms at these positions are lower than the protein-wide average frequency at a given ACE2 site (black line). (**D**)Systolic blood pressure divergence between rodents (red) and primates (blue) is correlated with differences in body size (*P*=0.001; phylogenetically independent least squares linear regression). (**E**) Rodent and bats species display higher ACE2 evolutionary rates relative to other species (aBSREL). (**F**) Phylogeny of jawed vertebrates used in the ancestral reconstruction of *ACE2* from the last common ancestor of rodents (RodAnc). (**G**) RodAnc ACE2 displays none of the rare sequence variants unique to the RBD binding domain of mice (**H**) Evolution of high ACE2 activity in the rodent ancestor. The C-terminus of mouse (M) or human (H) was used.

An alternative explanation is that these evolutionary valleys may not exist in rodents. Specifically, this trajectory may have been opened to mice by permissive mutations (*25*, *31*) that compensated for the subsequent introduction of rare variants that are strongly deleterious in dog and human ACE2 backgrounds. To understand if ancestral rodent ACE2 displayed high activity before the introduction of these mutations, we constructed a large jawed vertebrate *ACE2* phylogenetic dataset (Fig. 3F, S5) and reconstructed the ancestral rodent (RodAnc) *ACE2* using likelihood methods (*57*). Reconstructed RodAnc ACE2 had none of the rare sequence variants unique to the RBD binding domain of mice (Fig. 3G). This strongly suggests that these mutations appeared relatively recently in mouse ACE2 evolution. Since the C-terminus of vertebrate *ACE2* was too divergent to align, we constructed the full-length rodent ancestor (RodAnc) ACE2 coding-sequence with either mouse (-M) or human (-H) C-termini, and measured ACE2-specific hydrolysis activity. We found that RodAnc-M displayed high ACE2 hydrolysis activity indistinguishable from that of mouse, strongly suggesting that the rodent ancestor had high ACE2 hydrolysis rates (Fig. 3H). Moreover, the RodAnc with the human C-terminus displayed ACE2 activity significantly lower than mouse ACE2, but significantly higher than that of human, suggesting the C-terminus is necessary, but not sufficient to explain the high ACE2 activity seen in the RodAnc. Taken together, these results provide strong evidence that mutations outside the viral binding interface of ACE2 drove increases to ACE2 carboxypeptidase activity during rodent evolution. High ACE2 activity in the rodent ancestor may have therefore compensated for the subsequent introduction of rare variants that are strongly deleterious mutations in dog and human ACE2 backgrounds.

## Discussion

We have shown that six mutations abolishing SARS-CoV-2 RBD-ACE2 binding also individually disrupt ACE2 hydrolysis activity by up to 50-60%. This provides evidence that like other proteins, the catalytic activity of ACE2 can be mediated by the indirect structural effects of mutations to residues outside the active site (*27*, *47*, *50*). Although previous work demonstrated that inactivation of the ACE2 active site had no effect on SARS-CoV-1 and −2 binding, these mutations were directly within the active site far from the viral binding interface (*58*, *59*). The mutations investigated here are directly within the viral binding interface of SARS-CoV-2, and from species naturally immune to SARS-CoV-2 that still maintain ACE2 activity. The surprising finding that mutations abolishing binding also reduce ACE2 activity is supported by the fact that human ACE2 engineered to increase viral binding (T27Y, L79T, N330Y) also displays reduced ACE2 activity (*24*). Here we discuss the implications of pervasive epistasis in ACE2 for the evolution of ACE2 function, the SARS-CoV-2 host range and therapeutic design.

Our work demonstrates the essential importance of the ACE2 viral binding domain on catalytic function. The consequent constraints on the ACE2 amino acid sequence in humans and other primates strongly indicate that protective polymorphisms in the human population are unlikely, in contrast to examples such as CCR5, the co-receptor for HIV (*60*). Our analysis shows that allele frequencies for ACE2 missense polymorphisms at six sites necessary for SARS-CoV-2 binding are lower than the protein-wide average, reflecting the physiological importance of these positions in mediating ACE2 catalytic activity. Although potentially protective ACE2 polymorphisms have been predicted at other sites in the RBD-ACE2 interface (*61*, *62*), we show that statistically unlikely combinations of mutations are required to disrupt SARS-CoV-2 binding, each of which have severe effects on ACE2 activity. Given the importance of ACE2 enzymatic processing of Ang II to Ang-(1-7) in protection against pathogenic features of multiple cardiovascular and kidney diseases (*37*, *41*, *42*), it is possible that that the extant mutational combinations observed in nature (e.g. human, bat, dog, mouse, pangolin) may all represent alternative sequence ‘solutions’ (*34*, *52*) each uniquely required for an animal’s physiology, thereby representing fitness ‘peaks’ in the sequence landscape (*28*, *31*, *34*, *53*, *63*–*65*). Species variation in ACE2 hydrolysis rates we observed may reflect variation in renin activity levels, which are a major factor in determining circulating Ang-II levels in humans (*39*, *66*, *67*). It is therefore likely that a preponderance of selective processes has shaped ACE2 evolution, imposed by a combination of epistatic and physiological constraints on ACE2 sequence-function diversification, as discussed below.

Our results strongly imply that the host range of SARS-CoV-2 may be broader than both empirical and computational studies have suggested (*5*–*11*, *15*). In fact, our results suggest that the SARS-CoV-2 host range among non-primate mammals may be essentially unpredictable. This warrants extreme caution for public health efforts aiming to limit the risks of anthroponosis and zoonosis between humans and other species, which can form novel viral reservoirs where new mutations can evolve (*13*, *14*). This has already occurred on mink farms, where infection by humans led to the evolution of novel viral strains, which have since reinfected human populations (*11*). The demonstrated infection of minks and other non-felid carnivores by SARS-CoV-2 despite predictions to the contrary (*6*, *7*, *9*–*11*, *15*) challenges the utility of predicting infection risk from sequence homology to the human ACE2 viral binding interface (*15*–*17*). Our results reconcile this apparent conundrum by demonstrating that the wide host range of SARS-CoV-2 appears to originate from the structural effects of interacting amino acids within the ACE2 viral binding interface. As in other proteins, this intramolecular epistasis makes the sequence-phenotype relationship unpredictable (*32*–*36*); intramolecular epistasis has generated sequence diversity driving ACE2 functional diversification (*34*, *35*) but through similar conformations all bound by SARS-CoV-2 (*25*, *27*, *29*). Only in a few lineages has the epistasis of these sites been permitted to open trajectories to new conformations not recognized by SARS-CoV-2 (*46*). Importantly, these new conformations confer ACE2 activity equivalent (bat *vs* dog) or greater (mouse *vs* human) than species susceptible to SARS-CoV-2 infection. Why have these mutational combinations not evolved in other mammalian lineages? The ability to traverse evolutionary valleys is related to effective population sizes (*31*), which are negatively correlated with body size (*70*), suggesting that the ACE2 mutational combinations investigated here may have been generated by the effects of population size on mutational rates. However, given the physiological importance of ACE2 activity discussed above, the existence of functionally detrimental intermediates may also explain why these conformations did not evolve more frequently in evolutionary history (*28*, *31*, *34*, *53*, *63*, *64*). Our evidence suggests a confluence of ancient permissive mutations and permissive cardiovascular constraints likely opened the available pool of sequence variation, allowing functional *and* conformational diversification (*25*, *28*, *29*, *31*). The highly effective binding mechanism of SARS-CoV-2 (*21*–*23*, *71*) may therefore synergize with the epistasis of the ACE2 domain to grant the virus a wide-host range—one that is very difficult to predict.

Our study also has relevance for therapeutic design. Human recombinant soluble ACE2 (hrsACE2) is in active development as a strategy to neutralize SARS-CoV-2 by binding the viral spike protein (*67*, *78*). In human COVID-19 patients, the catalytic activity of hrsACE2 can help reduce angiotensin II levels as well as inflammation associated with COVID-19, likely through elevating Ang-(1-7) levels (*67*). hrsACE2 may therefore have the added benefit of minimizing the injury to multiple organs caused by viral-induced downregulation of ACE2 expression and renin-angiotensin hyperactivation (*40*, *73*, *79*–*82*). However, under normal physiological conditions, higher sACE2 plasma activity has been associated with increased pulmonary artery systolic pressure and ventricular systolic dysfunction (*83*). In these instances where patients have pre-existing conditions that can be exacerbated by increasing sACE2 activity it may be beneficial to make use of catalytically inactive hrsACE2 that binds RBD with similar efficiency (*58*). There is likely to be a spectrum of scenarios where variable levels of ACE2 activity is desirable, in which case recombinant non-human ACE2 can serve a key role. Our results also have implications for domesticated animals susceptible to infection, especially minks, for whom the SARS-CoV-2 is particularly transmissible (*10*, *11*). In lieu of a vaccine for minks and other non-human mammals important to agricultural production, genetic engineering efforts could feasibly introduce the six residues we observed to abolish viral binding but maintain activity in dog ACE2. This would remove a dangerous reservoir of SARS-CoV-2 wherein the virus can evolve and potentially challenge the efficacy of emerging vaccines.

## Acknowledgements

The authors thank the Johns Hopkins Genetic Resources Core Facility (RRID:SCR_018669) for DNA sequencing. This work was supported by research grants from the National Institutes of Health (EY022383 and EY022683; to E.J.D.) and Core Grant P30EY001765, Imaging and Microscopy Core Module.

## Data and materials Availability

All data is available in the manuscript or in the supplementary materials

## Supplementary Materials

### Materials and Methods

#### ACE2 and SARS-CoV-2 constructs

Wild-type hACE2 with a 1D4 (C9) C-terminal tag (TETSQVAPA) was a gift from Hyeryun Choe (Addgene plasmid # 1786; http://n2t.net/addgene:1786; RRID:Addgene_1786) (*84*). Site-directed mutagenesis primers were designed to induce single amino acid substitutions via PCR (QuickChange II, Agilent). Mouse ACE2 was cloned from cDNA synthesized from an RNA extraction of ileum tissue from a C57/BL6 mouse sacrificed in compliance with all regulations of the Johns Hopkins University Institutional Animal Care and Use committee. ACE2 coding sequences from dog (XM_014111329.2), pangolin (*Manis javanica*; XM_017650263.1), bat (*Rhinolophus sinicus*; XM_019746337.1) and the rodent ancestor were synthesized as gblocks (Integrated DNA technologies). All animal ACE2 inserts were cloned into a pEGFP-N1 vector, with eGFP C-terminal tag. For interspecies comparisons, hACE2 was also cloned into pEGFP-N1. The receptor binding domain of the SARS-CoV-2 S protein was obtained as a pcDNA3-SARS-CoV-2-S-RBD-Fc plasmid, as a gift from Erik Procko (Addgene plasmid # 141183; http://n2t.net/addgene:141183; RRID:Addgene_141183) (*85*). pcDNA3.1-SARS-Spike was a gift from Fang Li (Addgene plasmid # 145031; http://n2t.net/addgene:145031; RRID:Addgene_145031) (*21*). We used this plasmid as a template to clone the RBD of SARS-CoV-1 Spike protein into pcDNA3 with a Fc C-terminal tag. This RBD consisted of residues 318 to 510, as previously described (numbering as per AAP13441.1) (*86*).

#### Flow cytometry and immunoprecipitation

HEK293T cells were transfected with ACE2-1D4, or SARS-CoV-2 S protein RBD-Fc constructs using TransIT-X2 (Mirus). 24hrs after transfection the media of RBD-transfected cells was replaced with OptiPRO SFM (ThermoFisher). 72hrs after transfection, the media from RBD-transfected cells was concentrated using an Amicon Ultra-15 centrifugal filter unit with a 3000 kDa molecular weight cutoff. In parallel, ACE2-transfected cells were harvested and washed with PBS. For flow cytometry, 5.0 x 10^5^ cells were resuspended in 1mL incubation buffer [Dulbecco’s PBS containing 0.02% EDTA (Sigma-Aldrich), 50μg/mL DNase I (Worthington) and 5mM MgCl2] and incubated with 20 ug/mL of RBD-Fc for 30 minutes at room temperature as previously described (*19*). Cells were then washed with buffer (5% FBS, 0.1% Sodium Azide in PBS) and incubated with human ACE2 Alexa Fluro 647-conjugated antibody (1μg/10^6^ cell, #FAB9332R, R&D Systems), human IgG Fc PE-conjugated antibody (10μg/10^6^ cell, #FAB110P, R&D Systems) or human IgG Fc APC-conjugated antibody (10μg/10^6^ cell, #FAB110A, R&D Systems) for 30 minutes at room temperature. LSR II (BD bioscience) was used to collect the data and Flowjo (Flowjo, LLC.) was used for analysis, conducted as previously described (*19*). For immunoprecipitation, ACE2-transfected cells were lysed in a PBS buffer containing 1% CHASPO (Sigma-Aldrich), and incubated with Dynabeads Protein G (ThermoFisher) and 2μg of RBD-Fc concentrate. Dynabeads were washed with PBS (0.5% CHAPSO), and elutions were immunoblotted using antibodies against 1D4 (Abcam; ab5417) and Human IgG Fc (Abcam; ab97225).

#### Pseudovirus Assay

SARS-CoV-2 spike pseudotyped murine leukemia virus (MLV) were generated based on a published protocol (*87*). Briefly, HEK293T cells were transfected with three plasmids, which express MLV Gag and Pol, firefly luciferase reporter and SARS-CoV-2 S protein. 48 h after transfection, culture medium containing pseudotyped particles were centrifuged and then filtered through a sterile 0.45 μm pore-sized filter to remove cell debris. For transduction, HEK293T cells were first transfected with ACE2 orthologs using lipofectamine 3000. 24 hours later, 200μL of SARS-CoV-2 Spike pseudotyped MLV were added to each well and incubated for 2 days. The transduction efficiency was quantified by measuring the activity of luciferase using luciferase assay system (Promega) and GloMax 20/20 luminometer (Promega).

#### ACE2 hydrolysis assay

HEK293T cells were transfected with ACE2 constructs using TransIT-X2 (Mirus). 72hrs later cells were washed in PBS and lysed in ACE2 reaction buffer pH 6.5 (1M NaCl, 0.5% Triton X-100, 0.5mM ZnCl2, 75mM Tris-HCl). Cell lysates were diluted in reaction buffer to 0.5μg protein, incubated ± the ACE2-specific inhibitor 10μM DX600 (Cayman Chemical) for 20 minutes at room temperature, followed by addition of 100μM Mca-YVADAPK(Dnp) (R&D Systems) and incubation at 37°C. Fluorescence emission at 405 nm was measured at 10, 60, and 120 minutes using a microplate reader (BMG Labtech) after excitation at 320 nm. Hydrolysis rates were quantified as fluorescence units per minute, using the slope of fluorescence development between 10 and 120 minutes, as previously described (*43*). An ANOVA general linear model with fluorescence as the response variable, mutation as the factor, and assay time as the covariate was fit to fluorescence data generated in the ACE2 hydrolysis assay. Statistical differences in ACE2 hydrolysis rates were determined using a cross factor between the mutation factor and time covariate.

#### Phylogenetic Comparative Methods

ACE and ACE2 coding sequences from 107 species representing all major jawed vertebrate lineages were obtained from GenBank (Table S4), and aligned using PRANK followed by manual adjustment (*88*). This alignment was used to estimate a gene tree using PhyML 3.1 (*89*) (Fig. S5), with GTR selected using automatic model selection based on AIC values (*90*), and aLRT SH-like branch support. This ML tree was rooted using *ACE*, and recapitulated all major phylogenetic relationships (Fig. S5 (*91*)). Ancestral sequences and posterior probability distributions were inferred using the best fitting models in the codeml package of PAML 4.9 (Table S5) (*49*). For estimation of dN/dS values using random sites models in the codeml package (*49*) and FUBAR (*92*), ACE sequences were pruned from the dataset, non-mammalian *ACE2* was also pruned, and additional mammalian *ACE2* sequences added to increase sampling. This alignment represented 89 species (Table S1), and was used to infer a ML gene tree using IQ-Tree (Fig. S2), with the substitution model auto selected, followed by ultrafast bootstrap analysis and SH-aLRT branch tests (*93*). PAML random sites models were used to investigate evidence of positive selection (Table S2). To test for dN/dS differences amongst branches in the phylogeny, clade model D (CmD) (*94*) was used to analyze an *ACE2* alignment containing only those sites of interest in the ACE2 viral binding interface (Table S3). M3 with three site classes was used as the null model for CmD. All random sites and clade model PAML model pairs were statistically evaluated for significance by likelihood ratio tests (LRT) with a χ2 distribution.

We conducted a phylogenetic comparative analysis on systolic blood pressure dataset by combining bat data with mammalian data compiled from a previous study (*77*, *95*). We pruned this dataset to include only species with fossil calibrated divergence times (*91*) (Fig. S6). Body size values (grams) were obtained from the Ageing Genomic Resources AnAge database (*96*). We conducted a phylogenetically independent correlation analysis (least squares linear regression; Fig. S6) by using the PDTREE program of the PDAP module of MESQUITE (*97*, *98*) to calculate phylogenetically independent contrasts of log10 body mass (grams) systolic blood pressures (mmHg) as the dependent variable. Before independent contrasts were calculated, branch lengths reflecting divergence times were ln transformed to meet the assumptions of independent contrast analysis (*97*, *98*). To produce dN/dS estimates for branches in the untransformed phylogeny, we pruned the mammalian *ACE2* alignment to match the species represented in the blood pressure dataset, and subjected the phylogeny and the alignment to analysis by aBSREL(*99*).

**Supplementary Figure S1.**
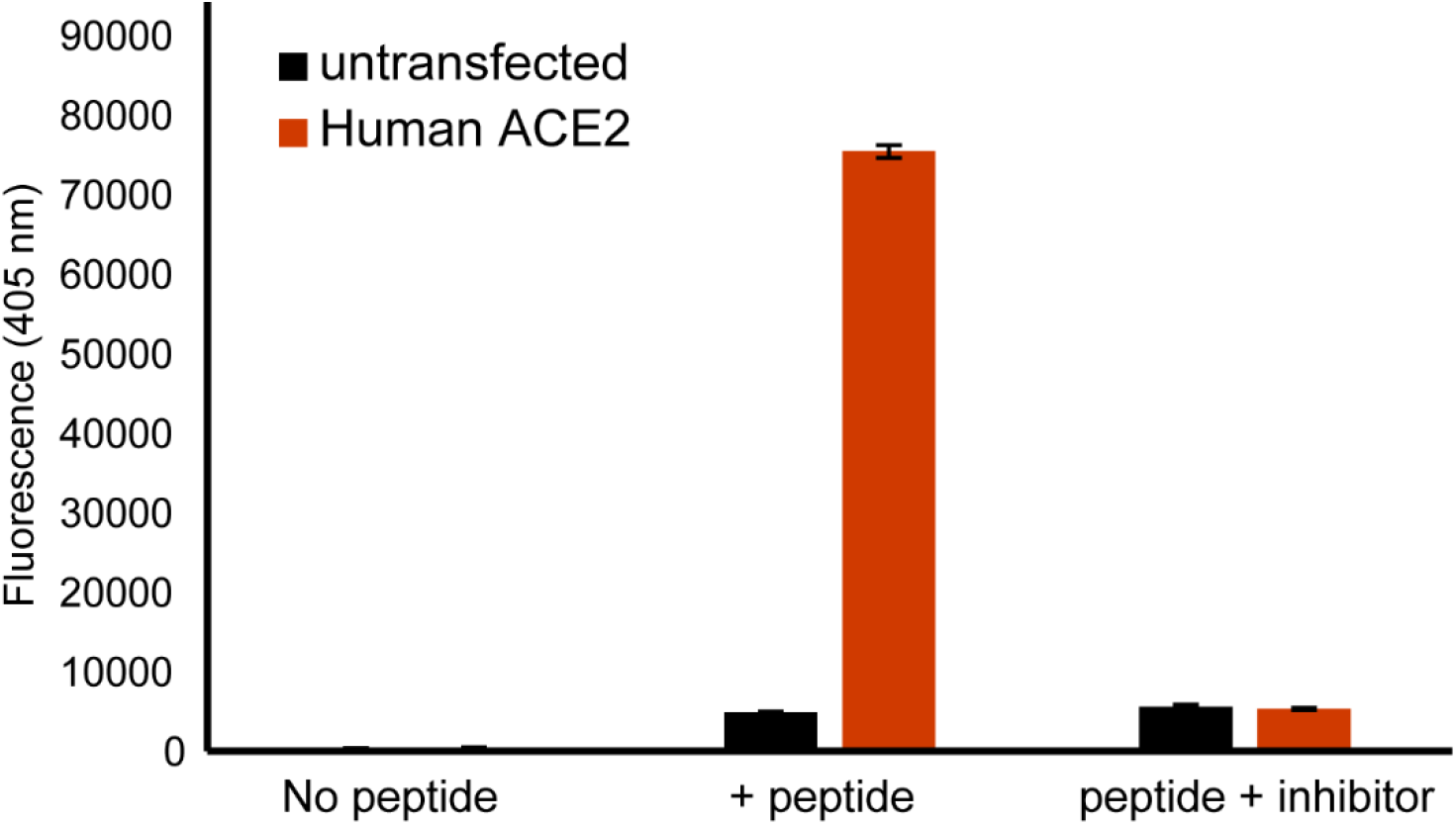
Recombinant ACE2 hydrolysis activity in solubilized transfected HEK293T cells. ACE2 hydrolysis activity was measured using a fluorogenic peptide substrate (Mca-YVADAPK(Dnp)-OH) incubated with lysates of HEK293T cells for 2 hours. Cells were either untransfected or were transfected with a human ACE2 construct. Incubation of lysate-peptide mixture with a ACE2-specific inhibitor (DX600) reduced fluorescence attributable to ACE2 hydrolysis activity.

**Supplementary Figure S2.**
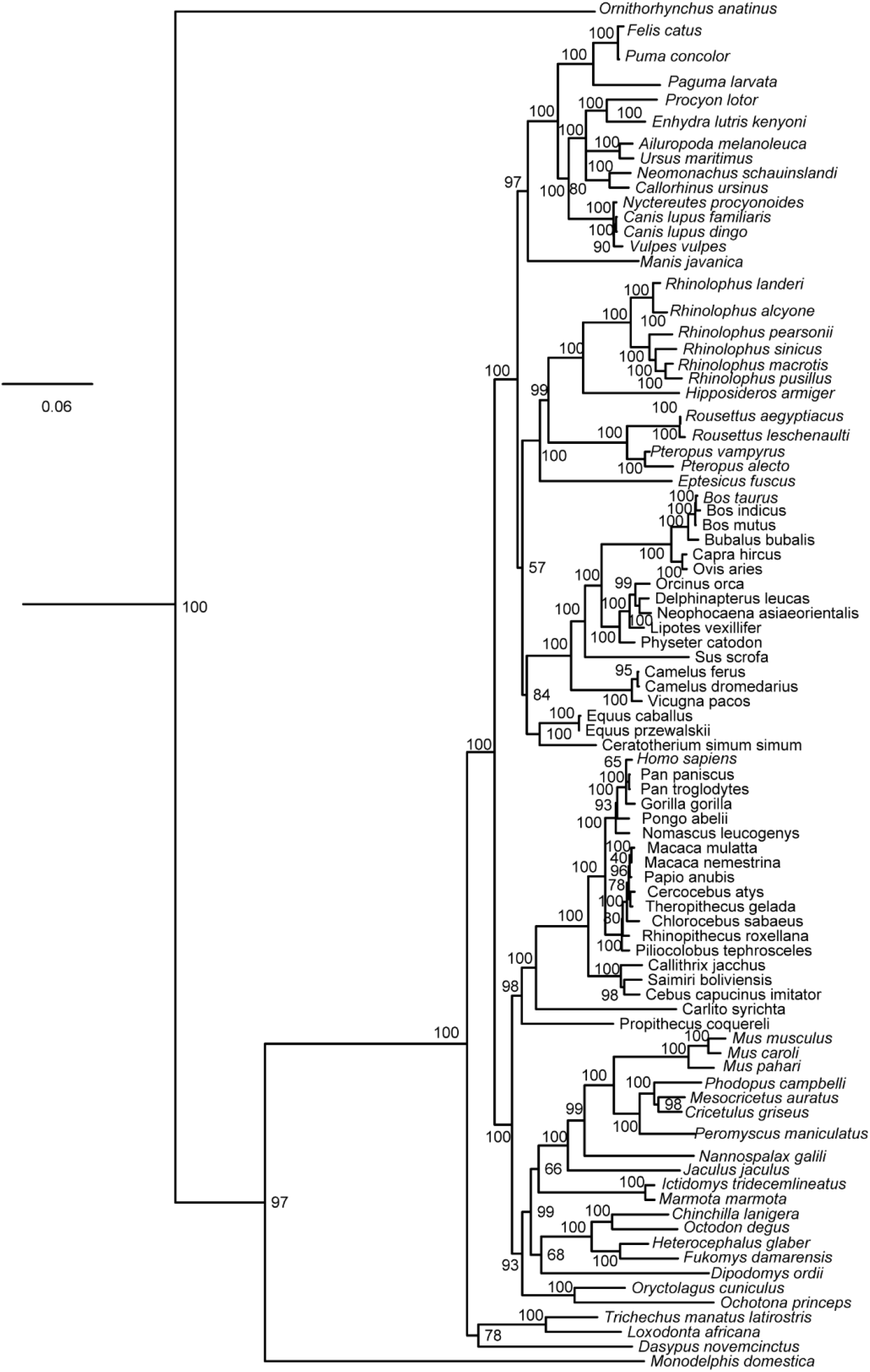
Maximum likelihood phylogeny used in PAML analyses. aLRT-SH like branch support values (PhyML) are shown

**Supplementary Figure S3.**
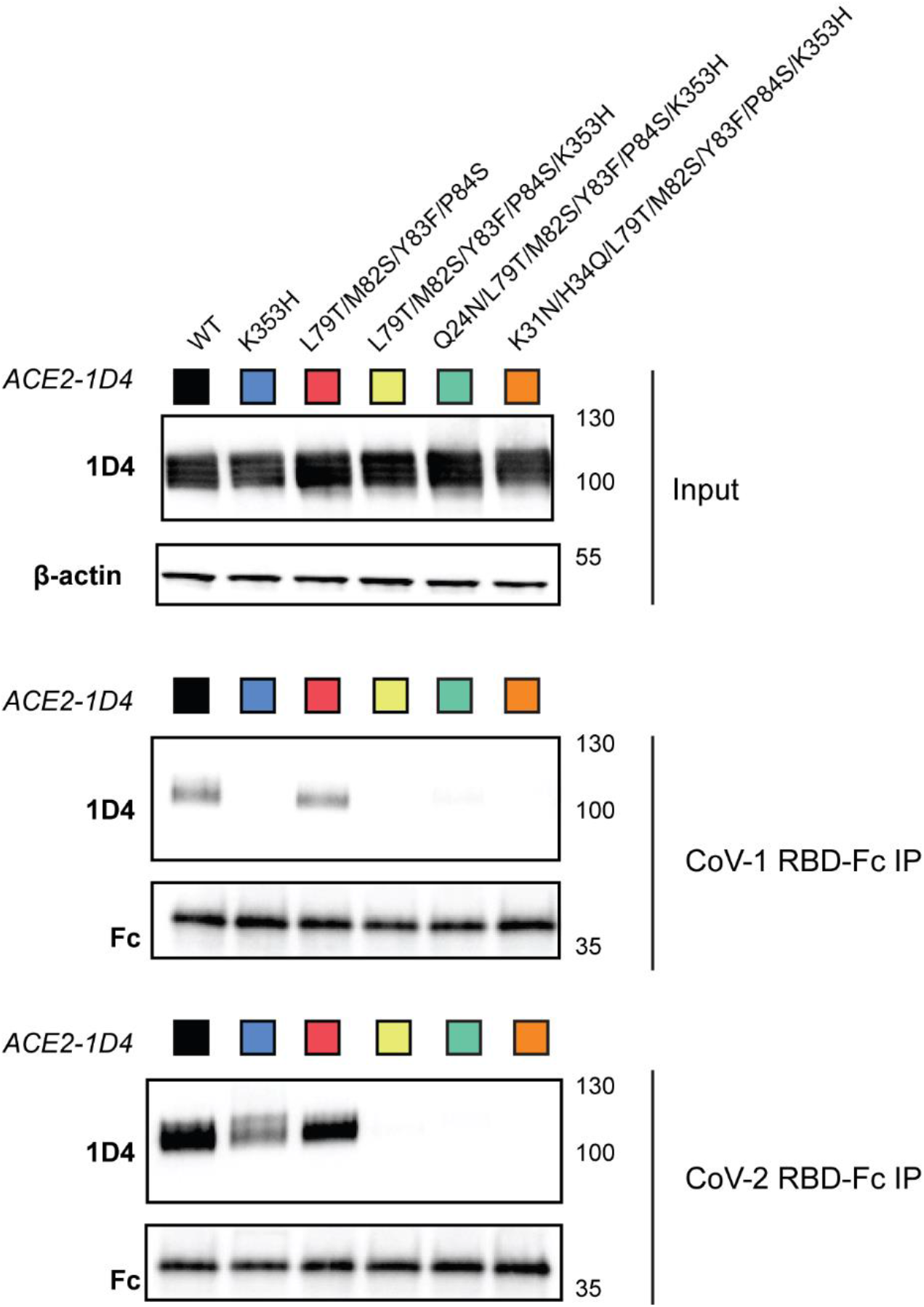
Targeted mutations to human ACE2 disrupt binding to the RBD of SARS-CoV-1 and SARS-CoV-2. Western blots of immunopreciptations and cell lysates of HEK293T cells co-transfected with an Fc-tagged SARS-CoV-2 S protein RBD, and 1D4-tagged (C9) human ACE2 construct.

**Supplementary Figure S4.**
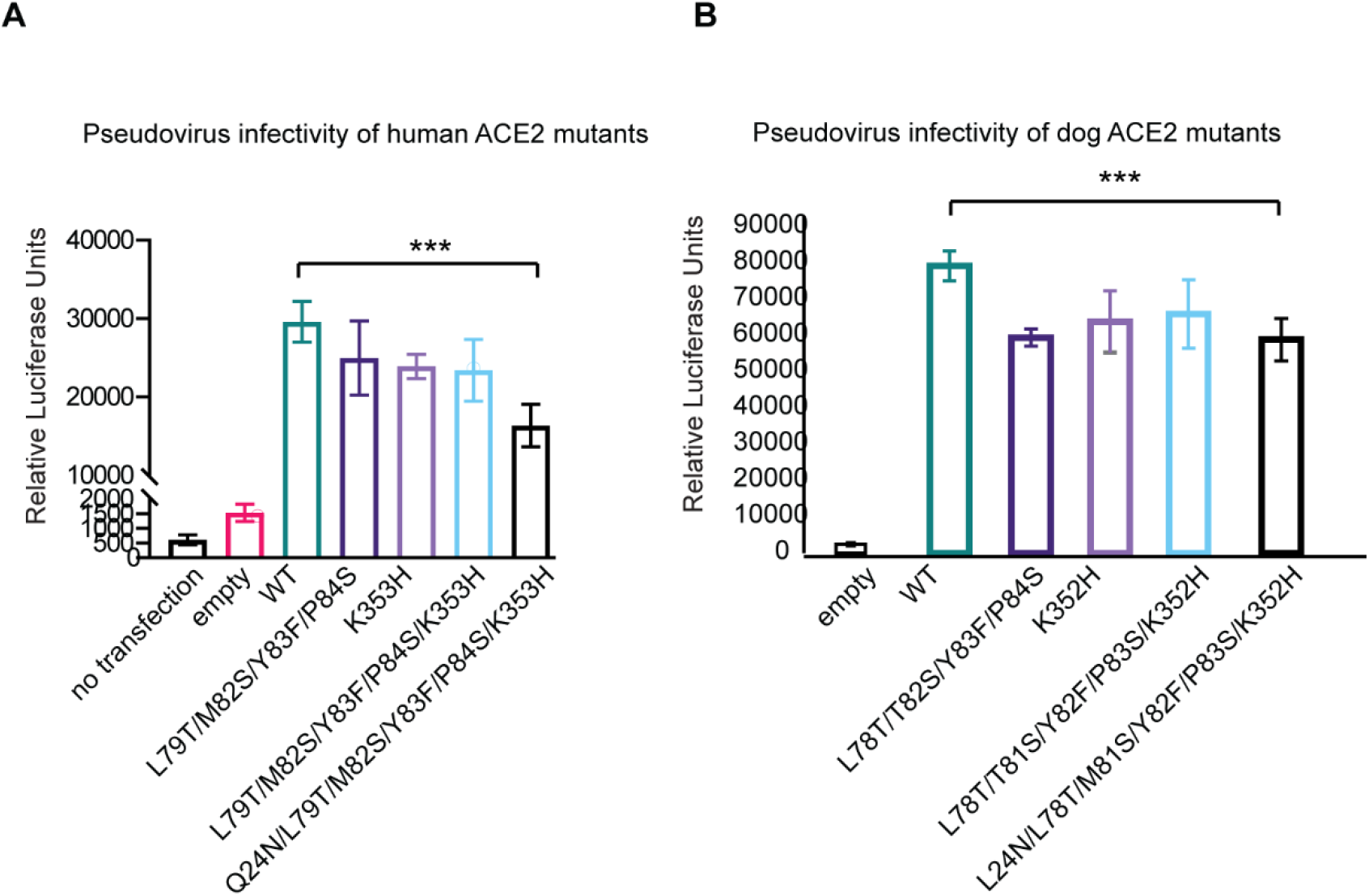
Effect of mouse substitutions on pseudovirus infection of cells expressing human and dog ACE2. Infection of HEK293T cells transfected with either (**A**) human or (**B**) dog ACE2, containing the indicated mutations. Cells were exposed to VSV-G pseudotyped with SARS-CoV-2 S protein, and cellular infection measured as a function of luciferase luminescence. (**C**) ACE2 hydrolysis activity of dog ACE2 with indicated mutations. ACE2 activity was measured as fluorescence units per minute.

**Supplementary Figure S5.**
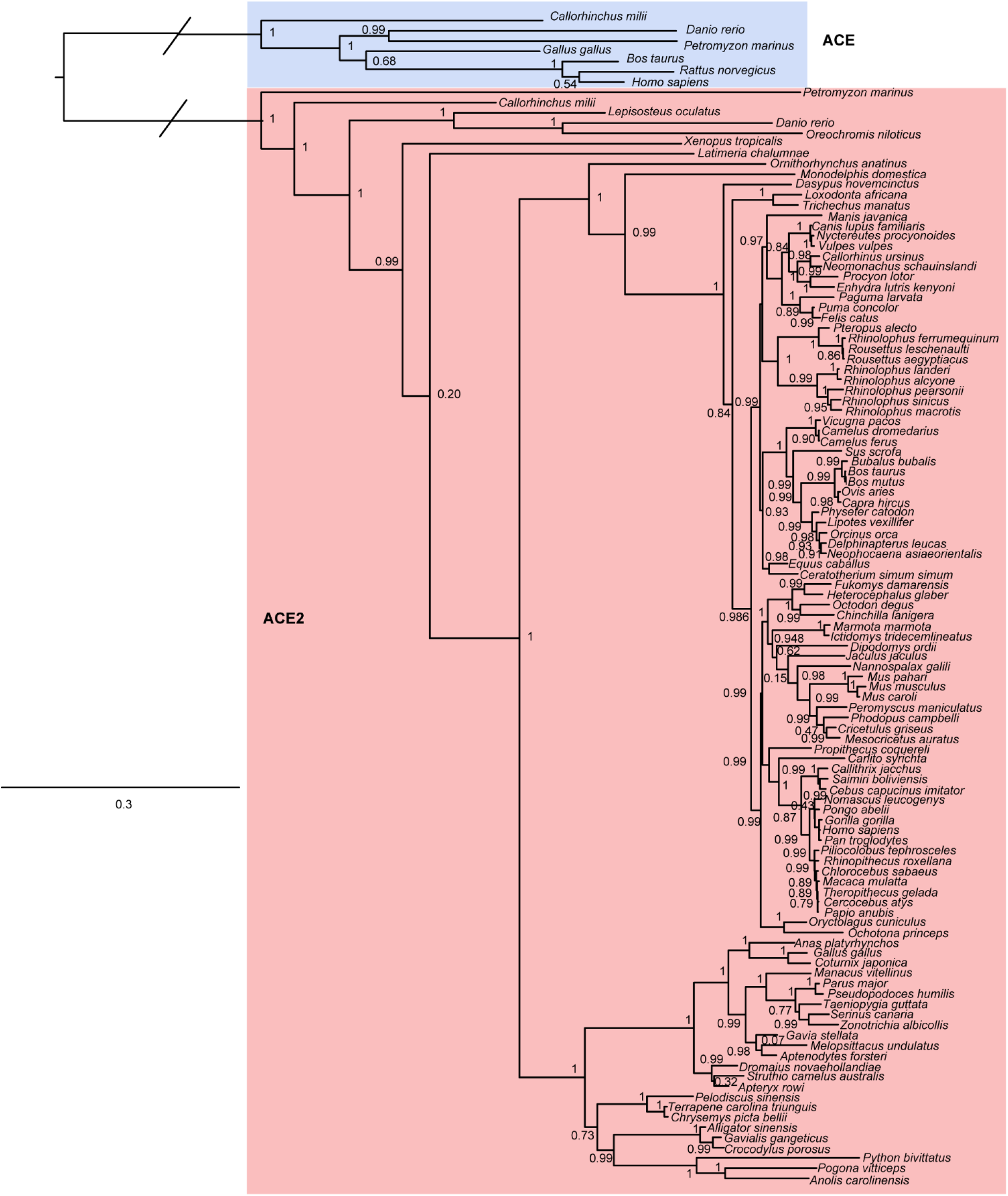
Maximum likelihood phylogeny used in ancestral reconstruction of Rodent ACE2. aLRT-SH like branch support values (IQ-Tree) are shown

**Supplementary Figure S6.**
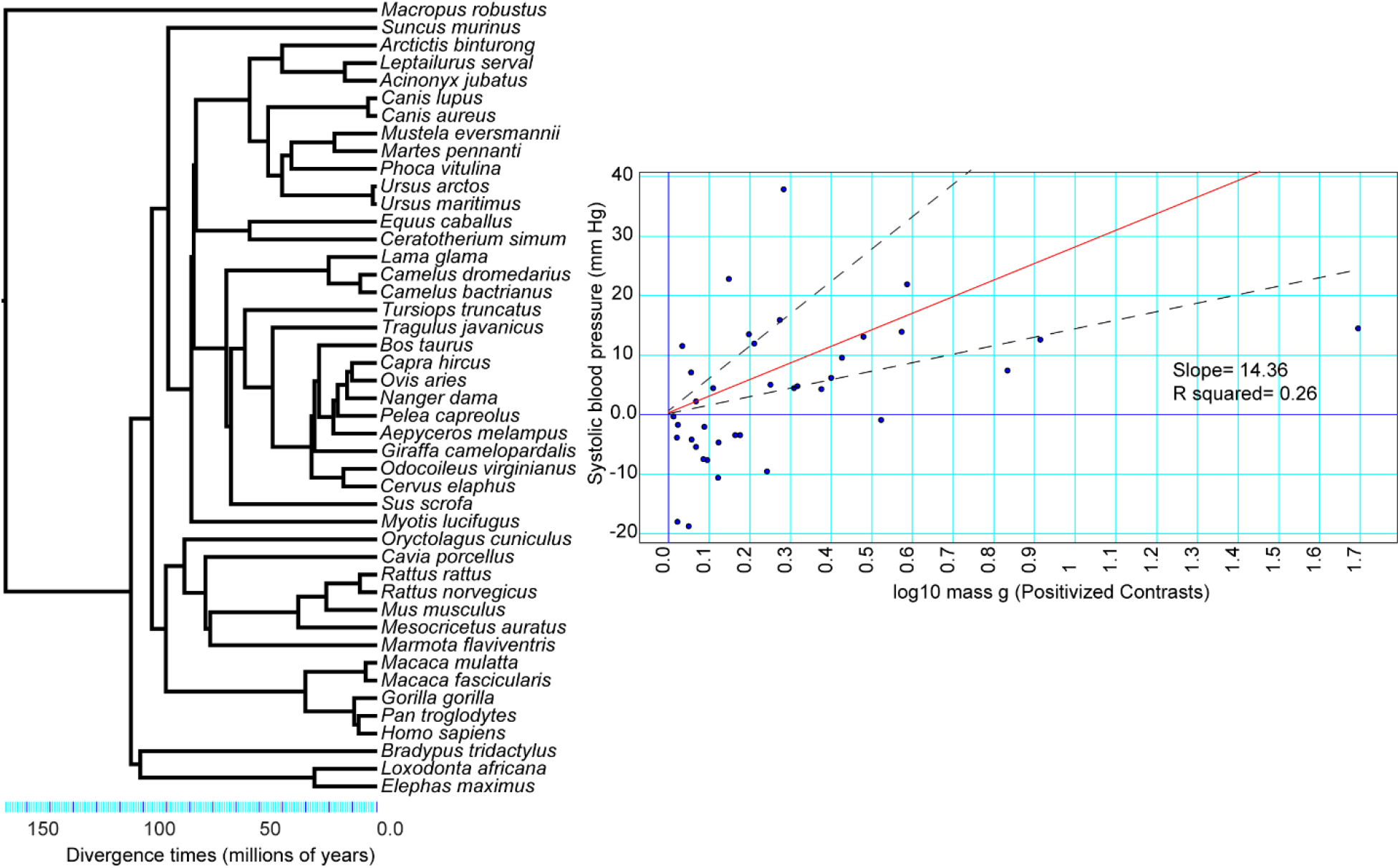
Species phylogeny and least squares linear regression using phylogenetically independent contrasts of systolic blood pressure and body mass

**Supplementary Table 1.**
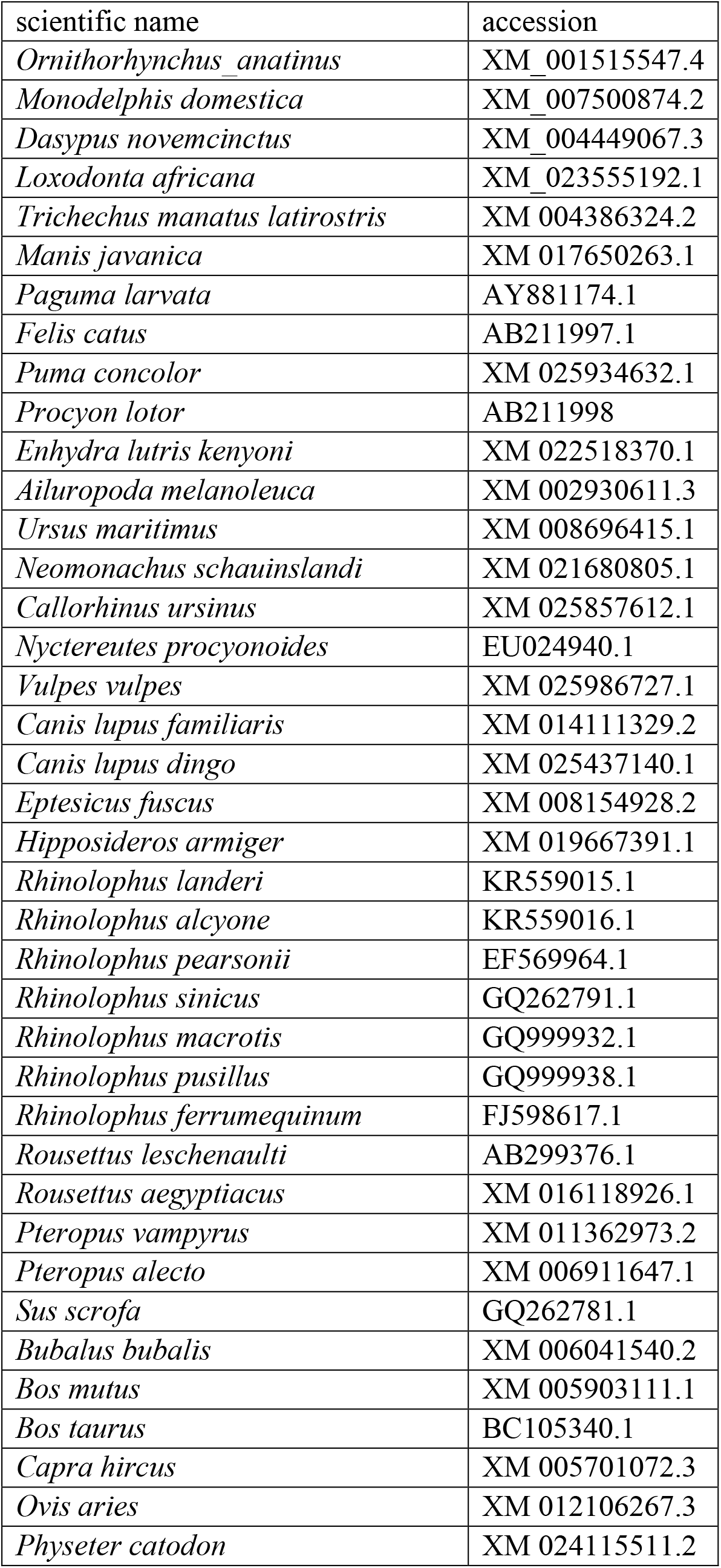
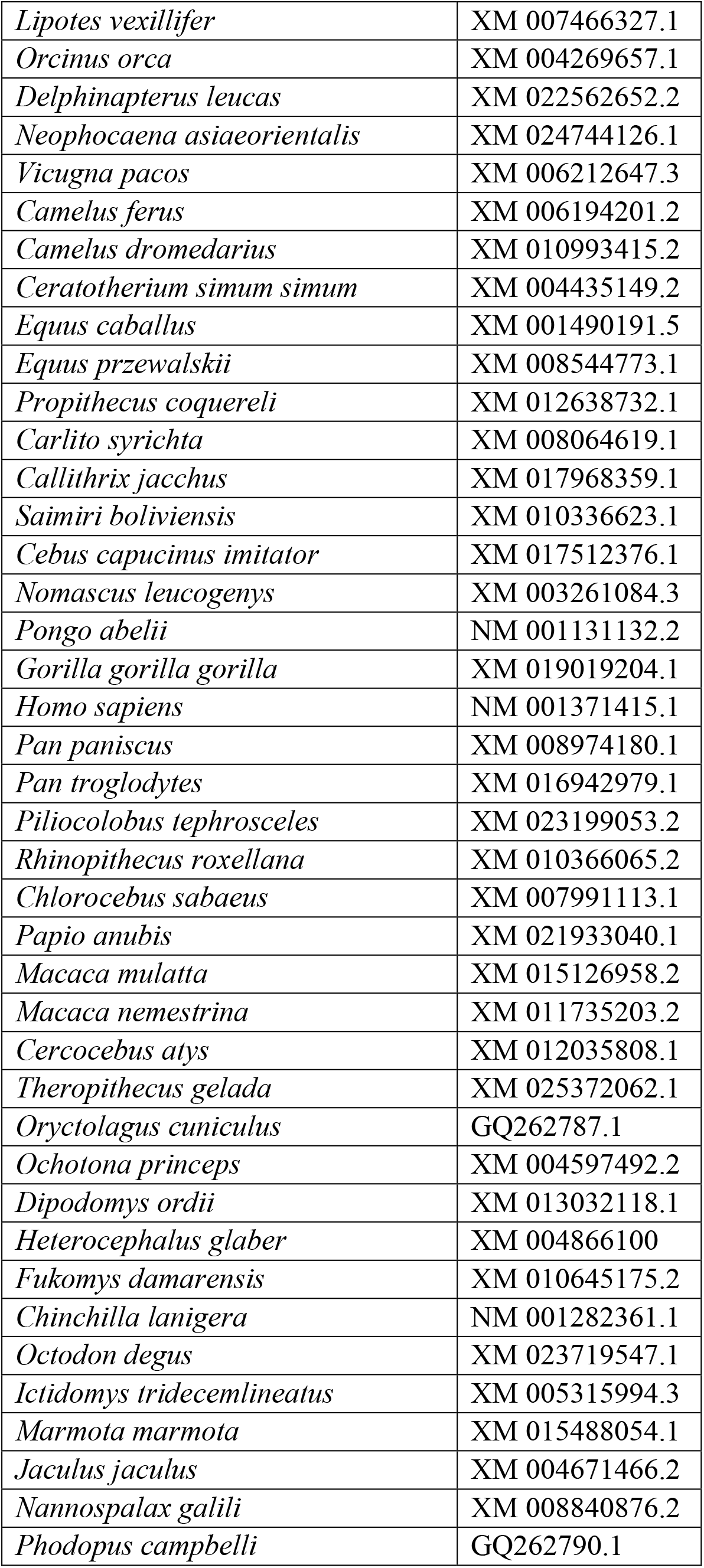
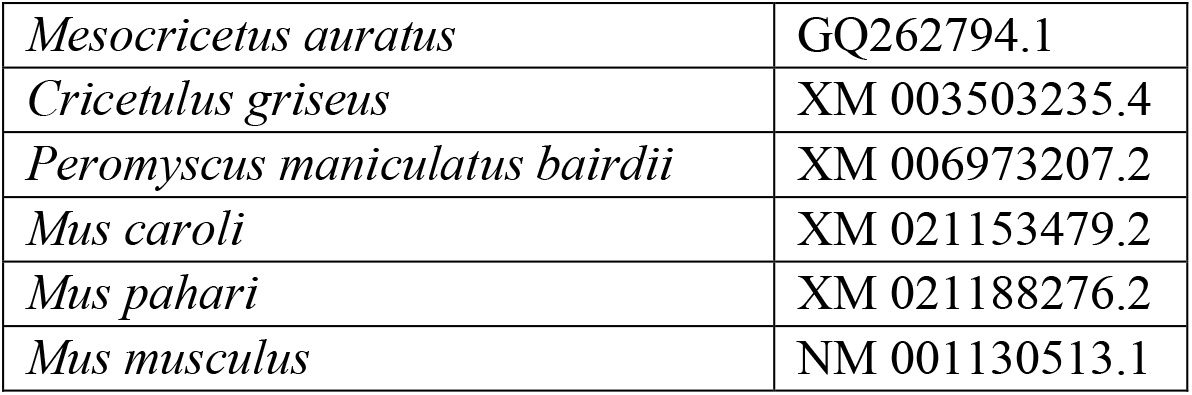
ACE2 accession numbers used in dN/dS estimates

**Supplementary Table 2.** Analyses of selection on Mammalian *ACE2* using PAML random sites models.

| Model | $\Delta\text{AIC}^1$ | $\ln L$ | Parameters<sup>2</sup> | | | Null | $p$ [df]<sup>3</sup> |
| --- | --- | --- | --- | --- | --- | --- | --- |
| | | | $\omega_0/p$ | $\omega_1/q$ | $\omega_2/\omega_p$ | | |
| M0 | 3746.8 | -36519.69 | 0.31 | - | - | N/A | - |
| M1a | 569.2 | -34929.93 | 0.08 (67%) | 1.00 (33%) | - | M0 | **0.000** [1] |
| M2a | 369.6 | -34828.09 | 0.08 (66%) | 1.00 (29%) | 2.39 (5%) | M1a | **0.000** [2] |
| M3 | 110.3 | -34697.44 | 0.04 (57%) | 0.48 (31%) | 1.51 (12%) | M2 | **0.000** [1] |
| M7 | 186.8 | -34738.71 | 0.24 | 0.55 | - | N/A | - |
| M8a | 141.3 | -34714.97 | 0.35 | 1.66 | 1.00 | N/A | - |
| **M8** | **0.0\*** | -34643.31 | 0.29 | 0.87 | 1.84 | M7 | **0.000** [2] |
|  |  |  |  |  |  | M8a | **0.000** [1] |

<sup>1</sup>All $\Delta\text{AIC}$ values are calculated from the lowest AIC model. The best fits are bolded with an asterisk (\*).
<sup>2</sup> $\omega$ values of each site class are shown for model M0-M3 ( $\omega_0$ – $\omega_2$ ) with the proportion of each site class in parentheses. For M7 and M8, the shape parameters, p and q, which describe the beta distribution are listed instead. In addition, the $\omega$ value for the positively selected site class ( $\omega_p$ , with the proportion of sites in parentheses) is shown for M8.
<sup>3</sup>Significant $p$ -values ( $\alpha \leq 0.05$ ) are bolded. Degrees of freedom are given in square brackets after the $p$ -values. Significance was determined through a likelihood-ratio test of null and alternative models, with reference to a $\chi^2$ distribution.
Abbreviations— $\ln L$ , ln Likelihood; $p$ , $p$ -value; N/A, not applicable.

**Supplementary Table 3.**
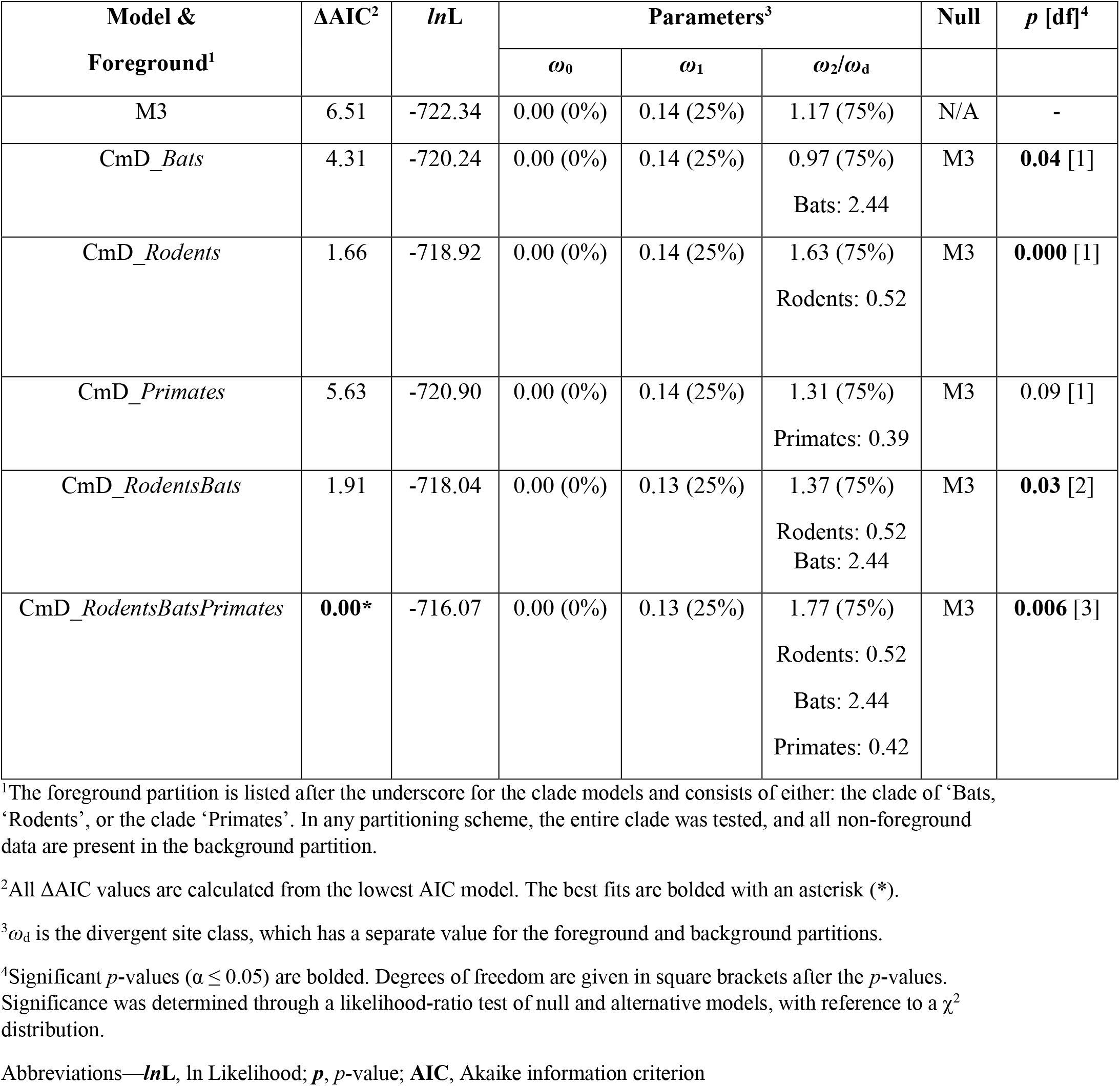
Results of Clade Model D (CmD) analyses of Mammalian *ACE2* under various partitions.

**Supplementary Table 4.**
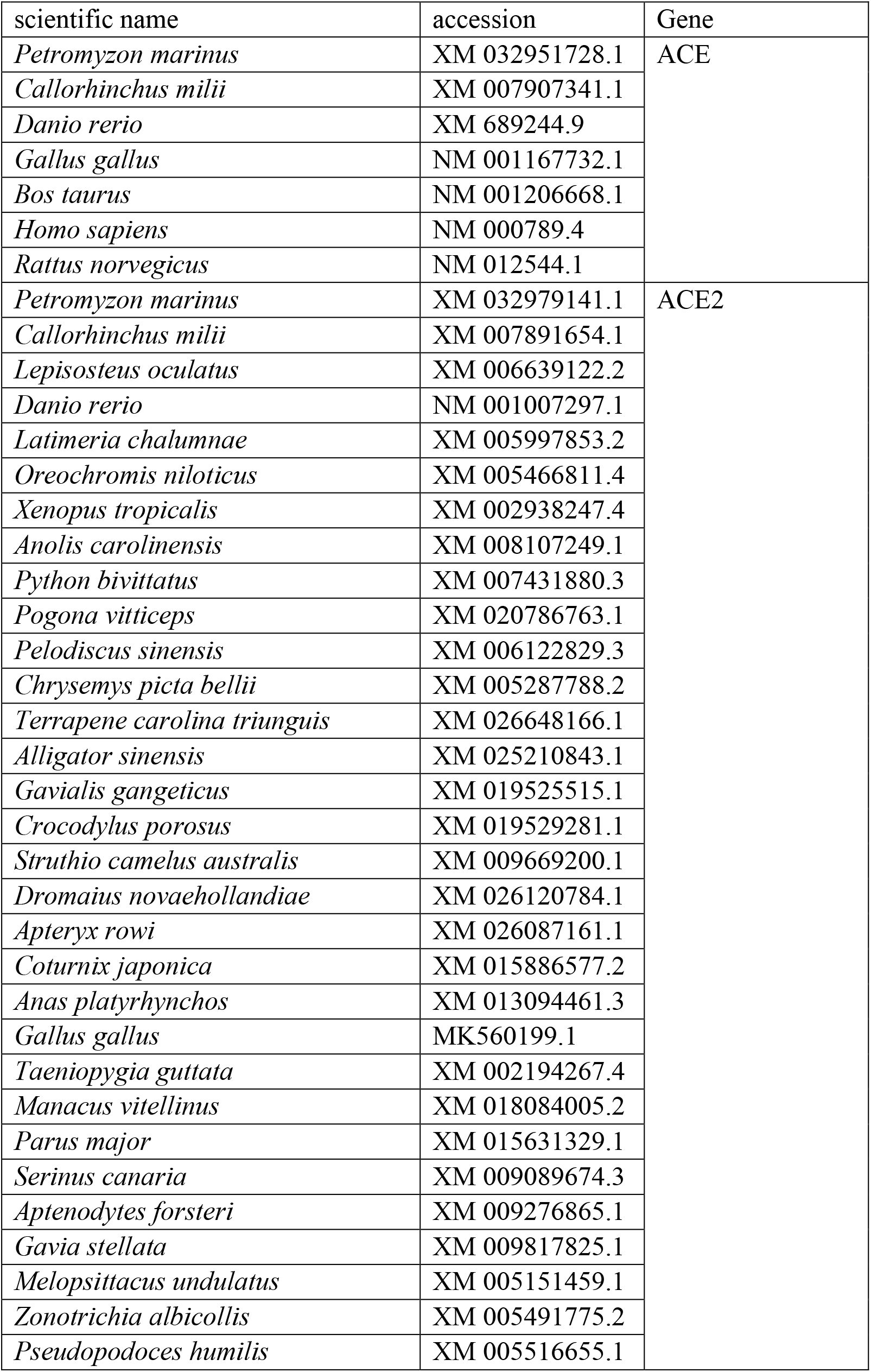
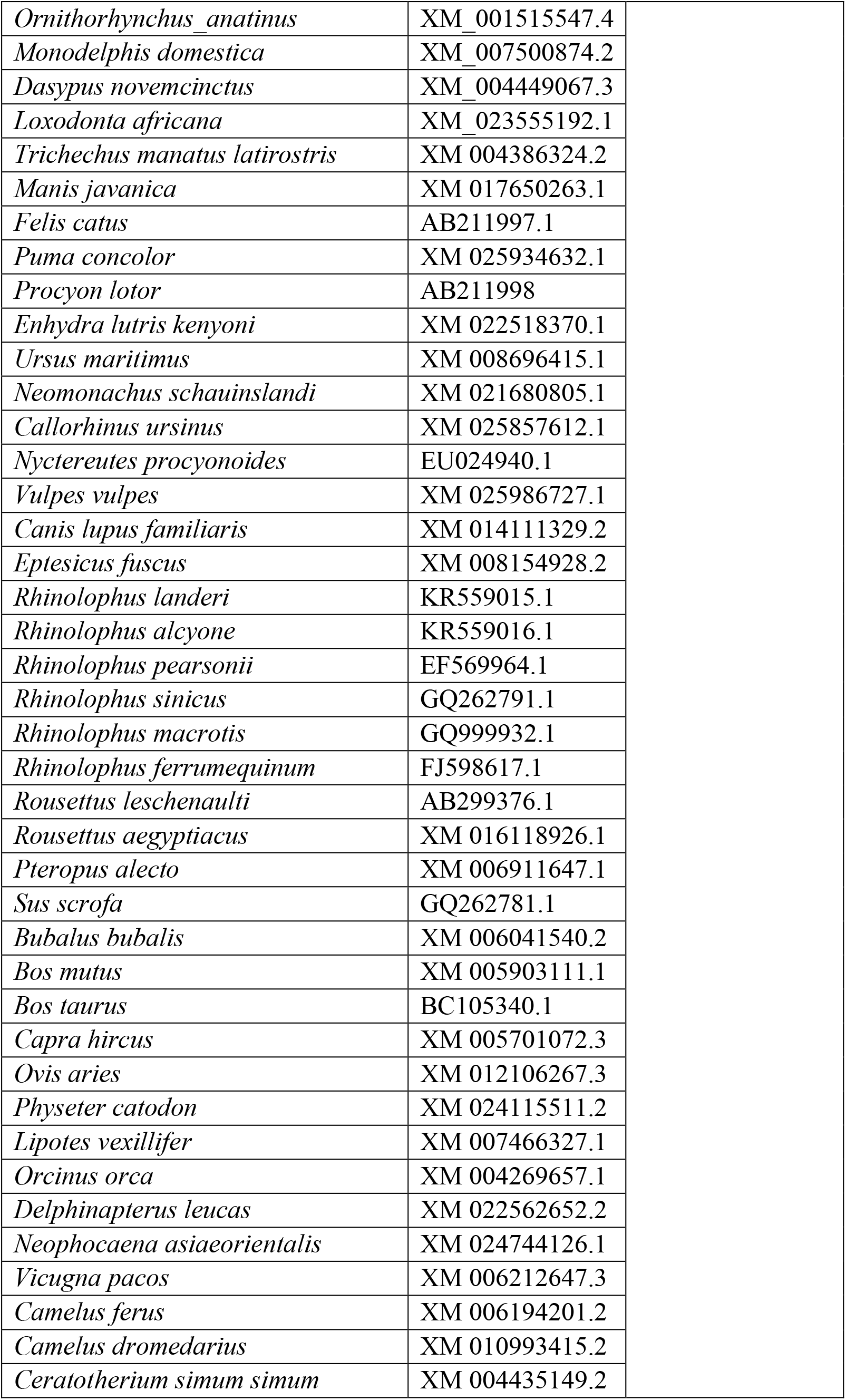
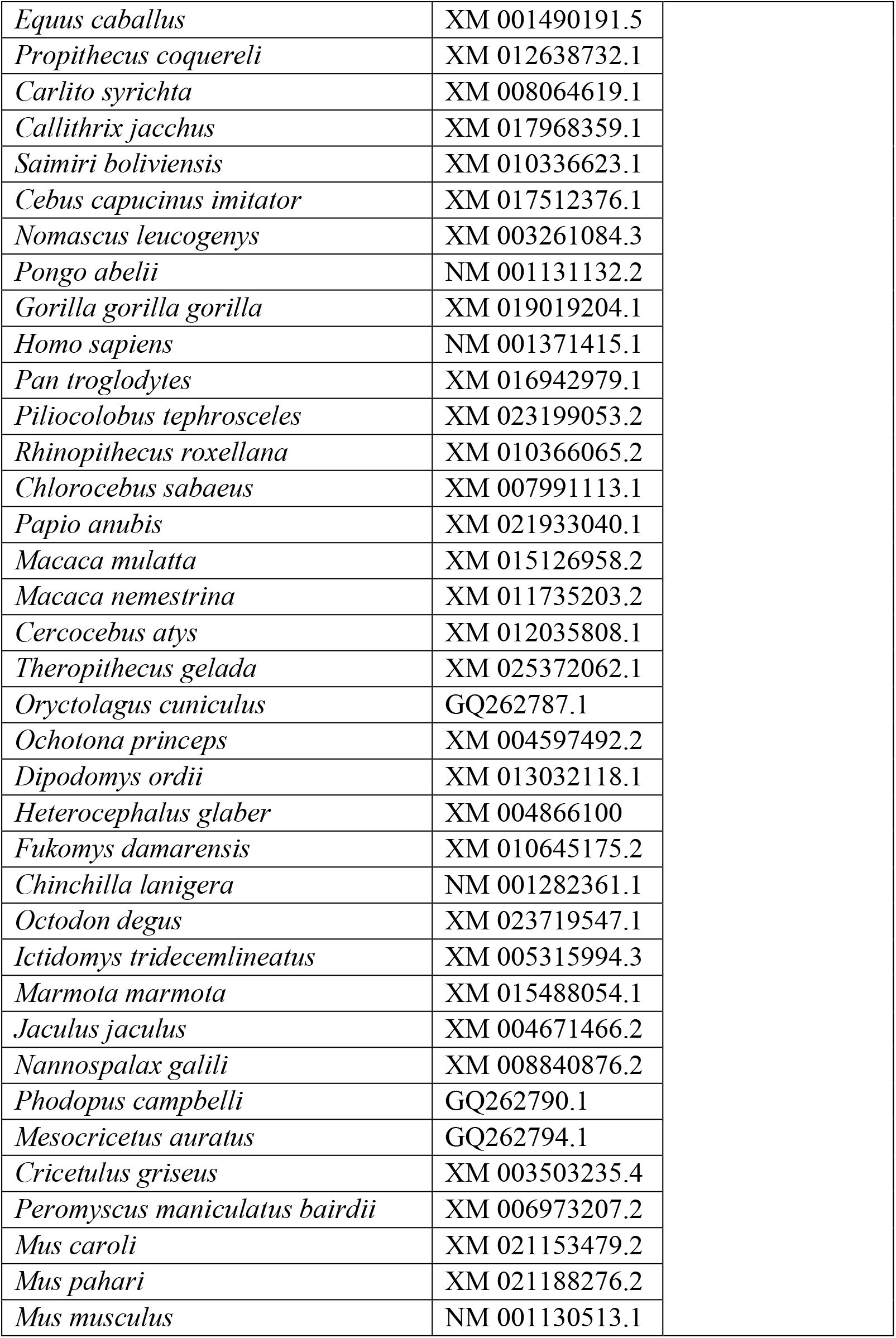
ACE and ACE2 accession numbers used in ancestral reconstruction

**Supplementary Table 5.** Results of random sites analyses of vertebrate *ACE2*, with the best fitting mode (M8) used for the ancestral reconstruction of mammalian ACE2.

| Model | ΔAIC | $\ln L$ | Parameters<sup>2</sup> | | | Null | $p$ [df]<sup>3</sup> |
| --- | --- | --- | --- | --- | --- | --- | --- |
| | | | $\omega_0/p$ | $\omega_1/q$ | $\omega_2/\omega_p$ | | |
| M3 | 231.6 | -63042.66 | 0.02 (42%) | 0.23 (33%) | 0.78 (25%) | N/A | - |
| M7 | 46.32 | -62953.03 | 0.35 | 0.84 | - | N/A | - |
| M8a | **53.04** | -62955.39 | 0.49 | 1.84 | 1 |  | - |
| **M8** | **0.0\*** | -62927.87 | 0.41 | 1.35 | 1.22 | M7 | **0.000** [2] |
|  |  |  |  |  |  | M8a | **0.000** [1] |

<sup>1</sup>All $\Delta$ AIC values are calculated from the lowest AIC model. The best fits are bolded with an asterisk (\*).
<sup>2</sup> $\omega$ values of each site class are shown for model M0-M3 ( $\omega_0$ – $\omega_2$ ) with the proportion of each site class in parentheses. For M7 and M8, the shape parameters, $p$ and $q$ , which describe the beta distribution are listed instead. In addition, the $\omega$ value for the positively selected site class ( $\omega_p$ , with the proportion of sites in parentheses) is shown for M8.
<sup>3</sup>Significant $p$ -values ( $\alpha \leq 0.05$ ) are bolded. Degrees of freedom are given in square brackets after the $p$ -values. Significance was determined through a likelihood-ratio test of null and alternative models, with reference to a $\chi^2$ distribution.
Abbreviations— $\ln L$ , ln Likelihood; $p$ , $p$ -value; N/A, not applicable.

